# Assessing the shared variation among high-dimensional data matrices: a modified version of the Procrustean correlation coefficient

**DOI:** 10.1101/842070

**Authors:** E. Coissac, C. Gonindard-Melodelima

## Abstract

**Motivation:** Molecular biology and ecology studies can produce high dimension data. Estimating correlations and shared variation between such data sets are an important step in disentangling the relationships between different elements of a biological system. Unfortunately, classical approaches are susceptible to producing falsely inferred correlations.

**Results:** Here we propose a corrected version of the Procrustean correlation coefficient that is robust to high dimensional data. This allows for a correct estimation of the shared variation between two data sets and the partial correlation coefficients between a set of matrix data.

**Availability:** The proposed corrected coefficients are implemented in the ProcMod R package available on CRAN. The git repository is hosted at https://git.metabarcoding.org/lecasofts/ProcMod

## 1 Introduction

Multidimensional data and even high-dimensional data, where the number of variables describing each sample is far larger than the sample count, is now routinely produced in functional genomics (*e.g.* transcriptomics, proteomics or metabolomics) and molecular ecology (*e.g.* DNA metabarcoding and metagenomics). Using a range of techniques, the same sample set can be described by several multidimensional data sets, each of them describing a different facet of the samples. This enables data analysis methods to evaluate mutual information shared by these different descriptions.

Correlative approaches are one of the simplest approaches to decipher pairwise relationships between multiple datasets. For a long time, several coefficients have been proposed to measure correlations between two matrices (for a comprehensive review see Ramsay *et al.*, 1984). However, when applied to high-dimensional data, these approaches suffer from over-fitting, resulting in high estimated correlations even for unrelated data sets. The creation of incorrect correlations from over-fitting consequently affects the biological interpretation of the analysis (Chariton *et al.*, 2010) can have downstream effects on the biological interpretation of a study. A number of modified matrix correlation coefficients have been proposed to address this issue. For example, the RV2 coefficient (Smilde *et al.*, 2009) corrects for over-fitting of the original RV coeffcient (Escoufier, 1973). Similarly, a modified version of the distance correlation coeffcient dCor (Székely *et al.*, 2007) proposed by SzéKely and Rizzo (2013) dCor has the advantage over the other correlation factors by considering by not being restricted to linear relationships.

Here we focus on the Procrustes correlation coefficient (RLs) proposed by Lingoes and Schönemann (1974) and by Gower (1971). Define *Trace*, a function summing the diagonal elements of a matrix. For an *n* × *p* real matrix **X** and a second *n* × *q* real matrix **Y** defining respectively two sets of *p* and *q* centered variables caracterizing *n* individuals, we define CovLs(**X**, **Y**) following Equation (1)

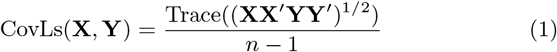

and VarLs(**X**) as CovLs(**X**, **X**). RLs can then be expressed as follow in Equation (2).

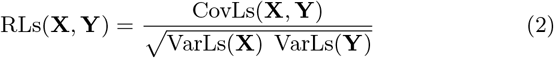

Considering CovLs(**X**, **Y**) and VarLs(**X**), respectively corresponding to the covariance of two matrices and the variance of a matrix, Equation (2) highlighting the analogy between RLs and Pearson’s correlation coefficient (R) (Bravais, 1844). When *p* = 1 and *q* = 1, RLs = |R|. Like the squared Pearson’s R, the squared RLs is an estimate of the amount of variation shared between the two datasets.

Procrustean analyses have been proposed as a good alternative to Mantel’s statistics for analyzing ecological data summarized by distance matrices (Peres-Neto and Jackson, 2001). In Procrustean analyze, distance matrices are projected into an orthogonal space using metric or non metric multidimensional scaling according to the geometrical properties of the used distances. Correlations can then be estimated on these projections.

## 2 Approach

RLs is part of the Procrustes framework that aims to superimpose a set of points with respect to another through three operations: a translation, a rotation, and a scaling. The optimal transfer matrix *Rot*_*X*→*Y*_ can be estimated from the singular value decomposition (SVD) of the covariance matrix *X′Y*. SVDs factorize any matrix as the product of three matrices (Equation 3).

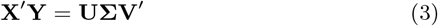

**U** and **V** are two rotation matrices which computes the transfer matrix to superimpose **X** on **Y** or reciproquely **Y** on **X**. All the elements of Σ except its diagonal are equal to zero. The diagonal elements are the singular values. Singular values are the extention of the eigenvalues to non-square matrices. CovLs(**X**, **Y**) can also be computed from singular values (Equation 4).

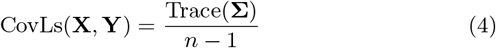

This expression actually illustrates that CovLs(**X**, **Y**) is the variance of the projections of **X** on **Y** or of the reciprocal projections. Therefore CovLs(**X**, **Y**) and RLs(**X**, **Y**) are always positive and rotation independant. Here we propose to partitionate Trace(**Σ**), the variation amount corresponding to CovLs(**X**, **Y**), in two components. The first corresponds to the actual shared information between **X** and **Y**. The second, corresponds to the over-fitting effect. It that can be estimated as the average variation shared by two random matrices of same structure as **X** and **Y** noted 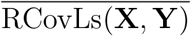. ICovLs(**X**, **Y**), the informative part of CovLs(**X**, **Y**), is computed using Equation (5).

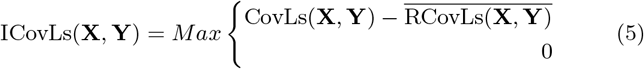

Similarly the informative counter-part of VarLs(**X**) is defined as IVarLs(**X**) = ICovLs(**X**, **X**), and IRLs(**X**, **Y**) the informative Procrustes correlation coefficient as defined in Equation (6.

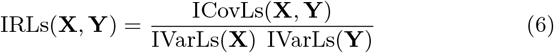

As in the case of RLs(**X**, **Y**), IRLs(**X**, **Y**) ∈ [0; 1], the 0 value corresponding to no correlation and the maximum value 1 reflects two strictly homothetic data sets.

The corollary of ICovLs(**X**, **Y**) and IVarLs(**X**) definitions is that ICovLs(**X**, **Y**) ≥ 0 and IVarLs(**X**) > 0. Therefore for *M* = {**M**_1_, **M**_2_, …, **M**_*k*_} a set of *k* matrices with the same number of rows, the informative covariance matrix **C** defined as **C**_*i,j*_ = ICovLs(**M**_*i*_, **M**_*j*_) is definite positive and symmetrical. This allows for defining the precision matrix **P** = **C**^−1^ and the related partial correlation coefficent matrix IRLs_*partial*_ using Equation (7)

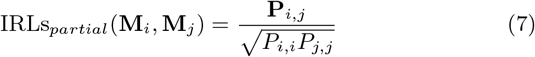

## 3 Methods

### 3.1 Monte-Carlo estimation of 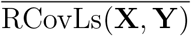

For every values of *p* and *q* including 1, 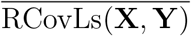 can be estimated using a series of *k* random matrices *RX* = {**RX**_1_, **RX**_2_, …, **RX**_*k*_} and *RY* = {**RY**_1_, **RY**_2_, …, **RY**_*k*_} where each **RX**_*i*_ and **RY**_*i*_ have the same structure as **X** and **Y**, respectively, in terms of number of columns and of the covariance matrix of these columns.

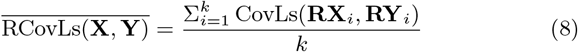

To estimate IVarLs(**X**), which is equal to ICovLs(**X**, **X**), 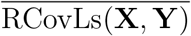 is estimated with two independent sets of random matrix *RX* and *RY*, both having the same structure than **X**.

#### Empirical assessment of 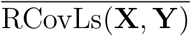

For two random vectors **x** and **y** of length *n*, the average coefficient of determination is 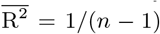. This value is independent of the distribution of the **x** and **y** values, but what about the independence of 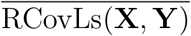 to the distributions of **X** and **Y** ? To test this independance and to assess the reasonnable randomization effort needed to estimate 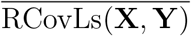, this value is estimated for four matrices **K**, **L**, **M**, **N** of *n* = 20 rows and 10, 20, 50, 100 columns, respectively. Values of the four matrices are drawn from a normal or an exponential distribution, with *k* ∈ {10, 100, 1000} randomizations tested to estimate 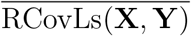 and the respective standard deviation 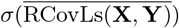. The VarLs of the generated matrices is equal to 1, therefore the estimated CovLs are equals to the RLs.

### 3.2 Simulating data for testing sensibility to over-fitting

To test overfitting, correlations were mesured between two random matrices of same dimensions. Each matrix is *n* × *p* with *n* = 20 and *p* ∈ [2, 50]. Each *p* variables were drawn from a centered and reduced normal distribution 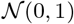. Eight correlation coefficients were tested: RLs the original Procrustes coefficient; IRLs this work; RV the original R for vector data (Robert and Escoufier, 1976); RV *adjM aye*, RV 2 and RV *adjGhaziri* three modified versions of RV (El Ghaziri and Qannari, 2015; Mayer *et al.*, 2011; Smilde *et al.*, 2009); dCor the original distance correlation coefficient (Székely *et al.*, 2007); and *dCor_ttest*, a modified version of dCor not sensible to overfitting (SzéKely and Rizzo, 2013). For each *p* value, 100 simulations were run. Computation of IRLs were estimated with 100 randomizations.

For *p* = 1, random vectors with *n* ∈ [3, 25] are generated. As above, data were drawn from 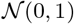 and *k* = 100 simulations which were run for each *n*. The original Pearson correlation coefficient *R* and the modified version IR are used to estimate correlation between both vectors.

### 3.3 Empirical assessment of the coefficient of determination

As in the case of the coefficient of determination (R^2^), RLs^2^ represents the part of shared variation between two matrices. Because of over-fitting in high-dimension data, RLs and therefore RLs^2^ are over-estimated.

#### Between two matrices

To test how the IRLs version of the coefficient of determination IRLs^2^ can perform to evaluate the shared variation, pairs of random matrices were produced for two values of *p* ∈ {10, 100} and *n* ∈ {10, 25}, and for several levels of shared variations ranging between 0.1 and 1 using 0.1 increments. For each combination of parameters, *k* = 100 simulations were run, and both RLs^2^ and IRLs^2^ were estimated using 100 randomizations.

#### Between two vectors

Coefficient of determination between two vectors also suffers from over-estimation when *n*, the number of considered points, is small. On average, for two random vectors of size *n*, 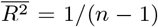. This random part of the shared variation inflates the observed shared variation even for non-random vectors. In the context of multiple linear regression, Theil (1958) proposed an adjusted version of the coefficient of determination (Equation 9), correcting for both the effect of the number of vectors (*p*) and the vector size (*n*).

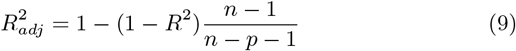

To evaluate the strength of that over-estimation and the relative effect of the correction proposed by Theil (1958) and by IRLs, pairs of random vectors were produced for *n* ∈ {10, 25}, and for several levels of shared variations ranging between 0.1 and 1 using 0.1 increments. For each combination of parameters, *k* = 100 simulations were run, and *R*^2^, 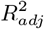 and IRLs^2^ were estimated using 100 randomizations.

#### Partial determination coefficients

To evaluate the capacity of partial determination coefficient 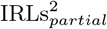 to distangle nested correlations, a set of correlated matrices were generated. To generate two random matrices **A**, **B**, sharing *w* ∈ [0, 1] part of variation, two independent random matrices **A** and **Δ** were generated such as VarLs(**A**) = 1 and VarLs(**Δ**) = 1.The **Δ**_*rot*_ matrix was computed as the alignment of **Δ** on **A** using the optimal Procrustes rotation. Then **B** is computed using equation 10:

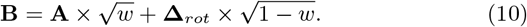

Following this method, four matrices **A**, **B**, **C**, and **D** of size *n* × *p* = 20 × 200 were generated where **A** shares 80% of variation with **B**, that shares 40% of variation with **C**, sharing 20% of variation with **D**. As illustrated in Figure 1, These direct correlations induce indirect ones spreading the total variation among each pair of matrices. The simulation was repeated 100 times, for every simutation 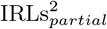 and 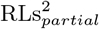 were estimated for each pair of matrices.

**Figure 1:**
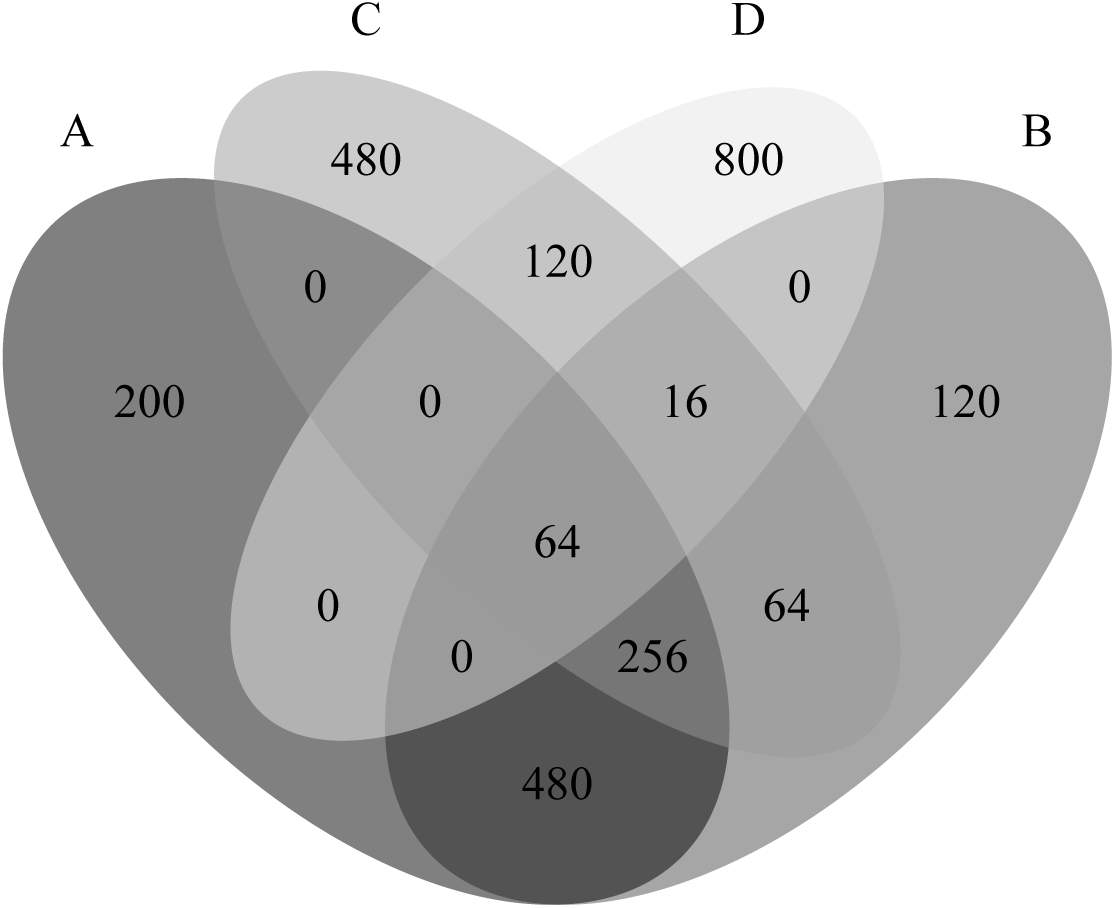
Theoretical distribution of the shared variation between the four matrices (**A**, **B**, **C**, **D**), expressed in permille.

### 3.4 Testing the significance of IRLs(X, Y)

The significance of IRLs(**X**, **Y**) can be tested using permutation test as defined in Jackson (1995) or Peres-Neto and Jackson (2001) and implemented respectively in the `protest` method of the vegan R package (Dixon, 2003) or the `procuste.rtest` method of the ADE4 R package Dray and Dufour (2007).

It is also possible to take advantage of the Monte-Carlo estimation of 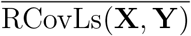 to test that ICovLs(**X**, **Y**) and therefore IRLs(**X**, **Y**) are greater than expected under random hypothesis. Over the *k* randomizations, *N*_>*CovLs*_ is estimated by counting when RCovLs(**X**, **Y**)_*k*_ > CovLs(**X**, **Y**). The P_value_ of the test then can be estimated following Equation (11).

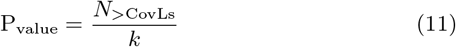

#### Empirical assessment of *α*-risk for the CovLs test

To empirically assess the *α*-risk of the Procrustes test based on the randomisations realized during the estimation of 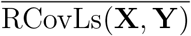, the distribution of *P*_*value*_ under *H*_0_ was compared to a uniform distribution between 0 and 1 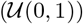. To estimate the empirical distribution, *k* = 1000 pairs of *n* × *p* random matrices with *n* = 20 and *p* ∈ {10, 20, 50} were simulated under the null hypothesis of independence. Significante of the Procrustes correlation between those matrices was tested using the three approaches: our proposed test (*CovLs.test*); the `protest` method of the vegan R package; the `procuste.rtest` method of the ADE4 R package. Conformance of the distribution of each set of *k P*_*value*_ to 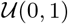 was assessed using the Cramer-Von Mises test (Csörgő and Faraway, 1996) implemented in the `cvm.test` function of the R package `goftest`.

#### Empirical power assessment for the *CovLs* test

To evaluate the relative the power of the three tests described above, pairs of two random matrices were produced for various *p* ∈ {10, 20, 50, 100}, *n* ∈ {10, 15, 20, 25} and two levels of shared variations R^2^ ∈ {0.05, 0.1}. For each combination of parameters, *k* = 1000 simulations were run. Each test were estimated based on 1000 randomizations for the *CovLs* test, or 1000 permutations for `protest` and `procuste.rtest`.

## 4 Results

### 4.1 Empirical assessment of 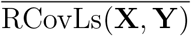

Two main parameters can influence the Monte Carlo estimation of 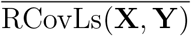: the distribution used to generate the random matrices, and *k* the number of random matrix pair. Two very different distribution are tested to regenerate the random matrices, the normal and the exponential distributions. The normal distribution is symmetric where the exponential is unsymmetrical with a high probability for small values and a long tail of large ones. Despite the use of these contrasted distributions, estimates of 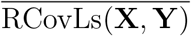 and of 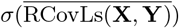 were identical if we assumed the normal distribution of the 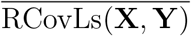 estimator and a 0.95 confidence interval of 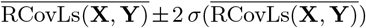 (Table 1).

**Table 1:**
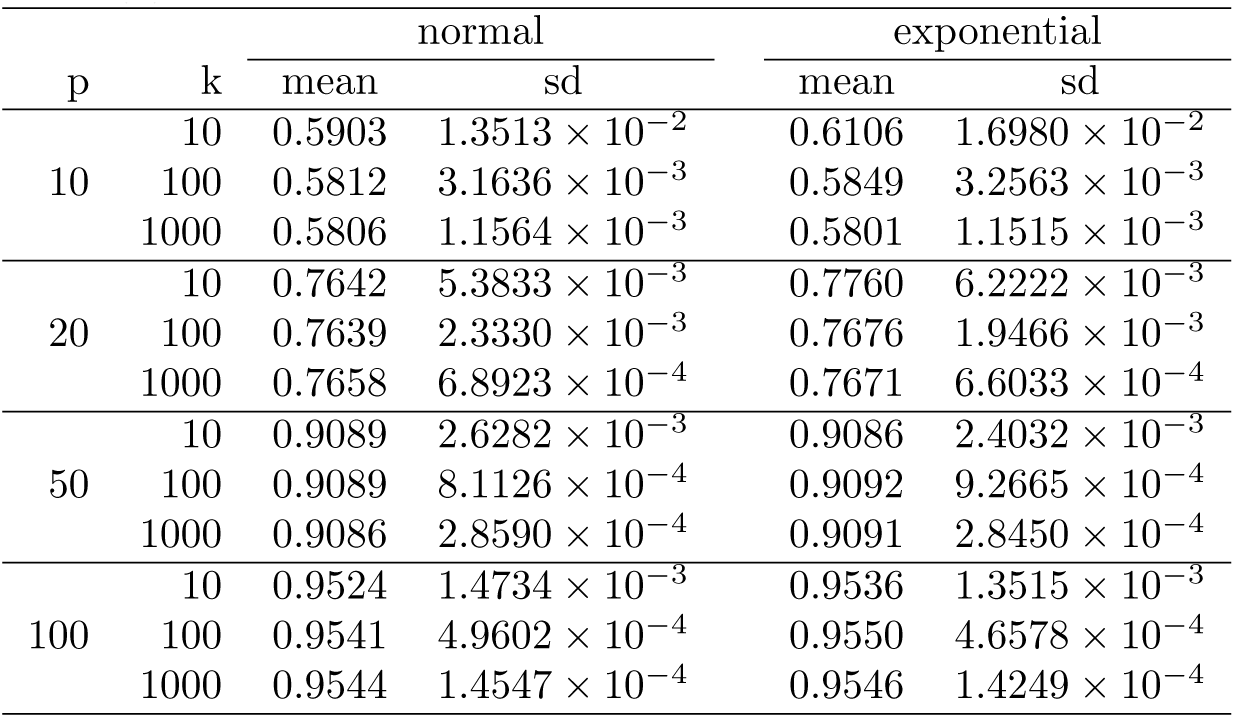
Estimation of 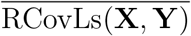 according to the number of random matrices (k) aligned.

| p | k | normal |  | exponential |  |
| --- | --- | --- | --- | --- | --- |
|  |  | mean | sd | mean | sd |
| 10 | 10 | 0.5903 | $1.3513 \times 10^{-2}$ | 0.6106 | $1.6980 \times 10^{-2}$ |
| | 100 | 0.5812 | $3.1636 \times 10^{-3}$ | 0.5849 | $3.2563 \times 10^{-3}$ |
| | 1000 | 0.5806 | $1.1564 \times 10^{-3}$ | 0.5801 | $1.1515 \times 10^{-3}$ |
| 20 | 10 | 0.7642 | $5.3833 \times 10^{-3}$ | 0.7760 | $6.2222 \times 10^{-3}$ |
| | 100 | 0.7639 | $2.3330 \times 10^{-3}$ | 0.7676 | $1.9466 \times 10^{-3}$ |
| | 1000 | 0.7658 | $6.8923 \times 10^{-4}$ | 0.7671 | $6.6033 \times 10^{-4}$ |
| 50 | 10 | 0.9089 | $2.6282 \times 10^{-3}$ | 0.9086 | $2.4032 \times 10^{-3}$ |
| | 100 | 0.9089 | $8.1126 \times 10^{-4}$ | 0.9092 | $9.2665 \times 10^{-4}$ |
| | 1000 | 0.9086 | $2.8590 \times 10^{-4}$ | 0.9091 | $2.8450 \times 10^{-4}$ |
| 100 | 10 | 0.9524 | $1.4734 \times 10^{-3}$ | 0.9536 | $1.3515 \times 10^{-3}$ |
| | 100 | 0.9541 | $4.9602 \times 10^{-4}$ | 0.9550 | $4.6578 \times 10^{-4}$ |
| | 1000 | 0.9544 | $1.4547 \times 10^{-4}$ | 0.9546 | $1.4249 \times 10^{-4}$ |

### 4.2 Relative susceptibility of *IRLs*(*X, Y*) to over-fitting

*RLs*, like *RV* and *dCor*, is susceptible to over-fitting which increases when *n* decreases, and *p* or *q* increase. Because *RV* is more comparable to *R*^2^, when *RLs* and *dCor* are more comparable to *R*, *RV* values increase more slowly than *RLs* and *dCor* values with *p* (Figure 2A). As expected *IRLs* values for non-correlated matrices are close to 0 regardless of *p* (Figure 2A). The same correction of the overfitting effect can be observed for vectors (Figure 2B)

**Figure 2:**
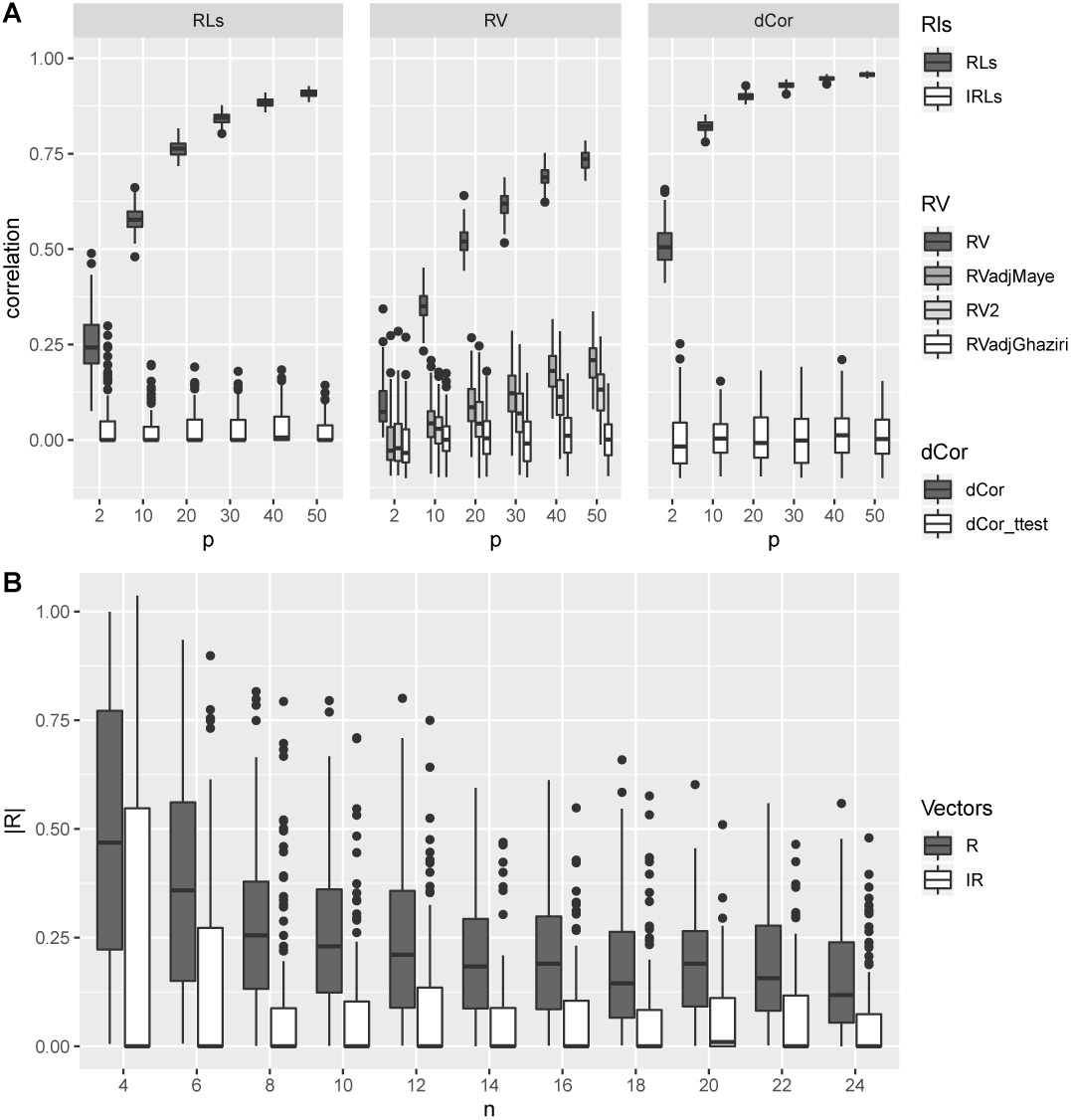
Susceptibility to overfitting for various correlation coefficients. A) Both simulated data sets are matrices of size (*n* × *p*) with *p* > 1. B) Correlated data sets are vectors (*p* = 1) with a various number of individuals *n* (vector length). For both A & B, 100 simulations were run for each combination of parameters.

### 4.3 Evaluating the shared variation

#### Between two matrices

For two matrices, our proposed corrected version of RLs^2^ (IRLs^2^) provided a good estimate of the shared variation and was robust to the phenomenon of over-fitting (Figure 3). Only a small over estimation was observed when *p* = 10 for the lowest values of the simulated shared variations (≤ 0.4).

**Figure 3:**
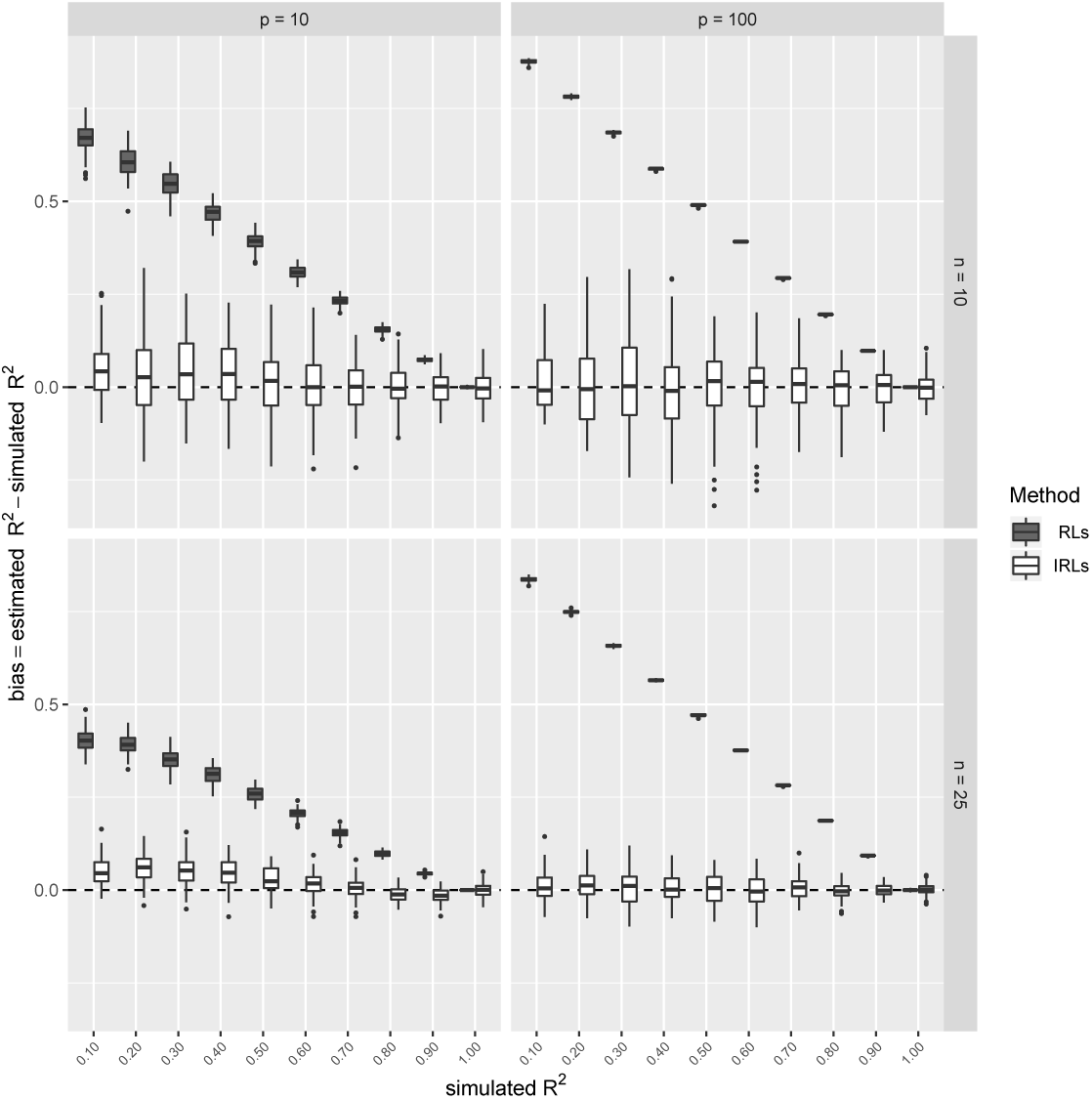
Shared variation (*R*^2^) between two matrices has measured using both the corrected (IRLs) and the original (RLs) versions of the Procrustean correlation coefficient. A gradient of *R*^2^ was simulated for two population sizes (*n* ∈ {10, 25}) and two numbers of descriptive variables (*p* ∈ {10, 100}). The distribution of differences between the observed and the simulated shared variation is plotted for each condition. The black dashed line corresponds to a perfect match where measured *R*^2^ equals the simulated one.

#### Between two vectors

Vectors can be considered as a single column matrix, and the efficiency of IRLs^2^ to estimate shared variation between matrices can also be used to estimate shared variation between two vectors. Other formulas have been already proposed to better estimate shared variation between vectors in the context of linear models. Among them the one presented in Equation 9, is the most often used and is the one implemented in R linear model summary function. On simulated data, IRLs^2^ performs better than the simple *R*^2^ and its modified version 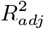 commonly used (Figure 4). Whatever the estimator the bias decrease with the simulated shared variation. Nevertheless for every tested cases the median of the bias observed is smaller than with both other estimators, even if classical estimators well perfom for large values of shared variation.

**Figure 4:**
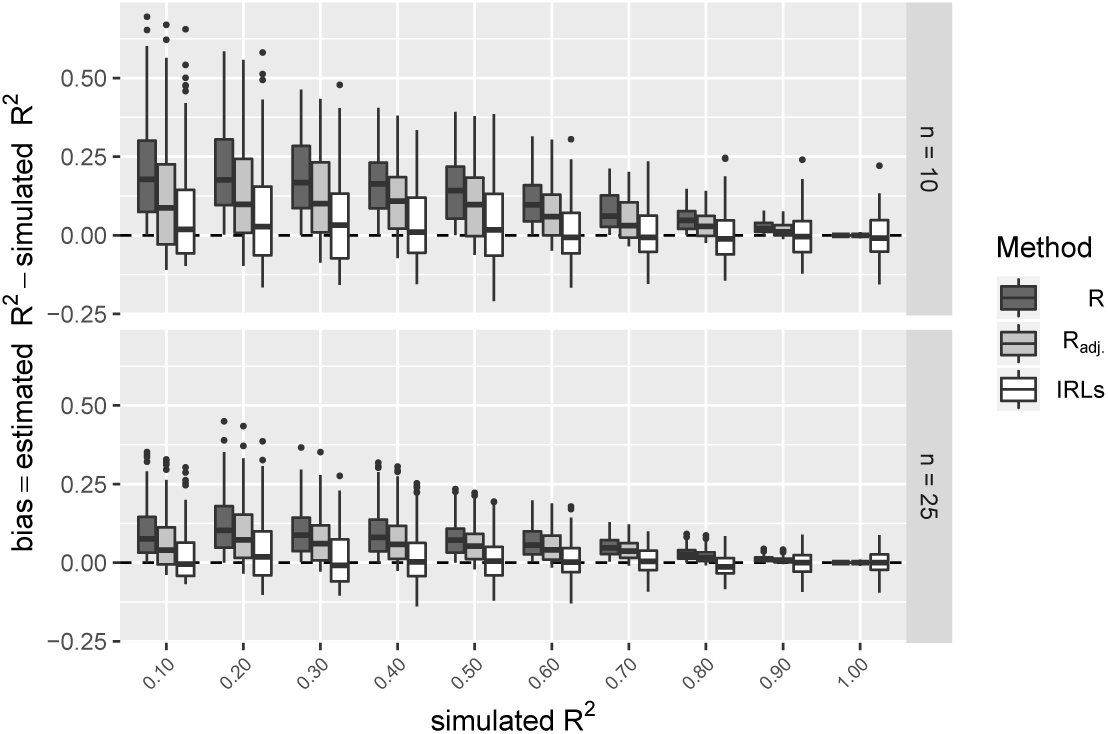
Shared variation between two vectors is measured using the classical R^2^, its adjusted version 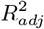 and (IRLs^2^). A gradient of shared variation is simulated for two vector sizes (*n* ∈ {10, 25}). The black dashed line corresponds to a perfect match where measured *R*^2^ equals the simulated one.

#### Partial coefficient of determination

The simulated correlation network between the four matrices **A**, **B**, **C**, **D** induced moreover the direct simulated correlation a network of indirect correlation and therefore shared variances (Figure 1). In such system, the interest of partial correlation coefficients and their associated partial determination coefficients is to measure correlation between a pair of variable without accounting for the part of that correlation which is explained by other variables, hence extracting the pure correlation between these two matrices. From Figure 1, the expected partial shared variation between **A** and **B** is 480/(200 + 480) = 0.706; between **B** and **C**, 64/(480 +120) = 0.107; and between **C** and **D** 120/800 = 0.150. All other partial coefficient are expected to be equal to 0. The effect of the correction introduced in IRLs is clearly weaker and on the partial coefficient of determination (Figure 5) than on the full coefficient of determination (Figure 3). The spurious random correlations, constituting the over-fitting effect, is distributed over all the pair of matrices **A**, **B**, **C**,**D**.

**Figure 5:**
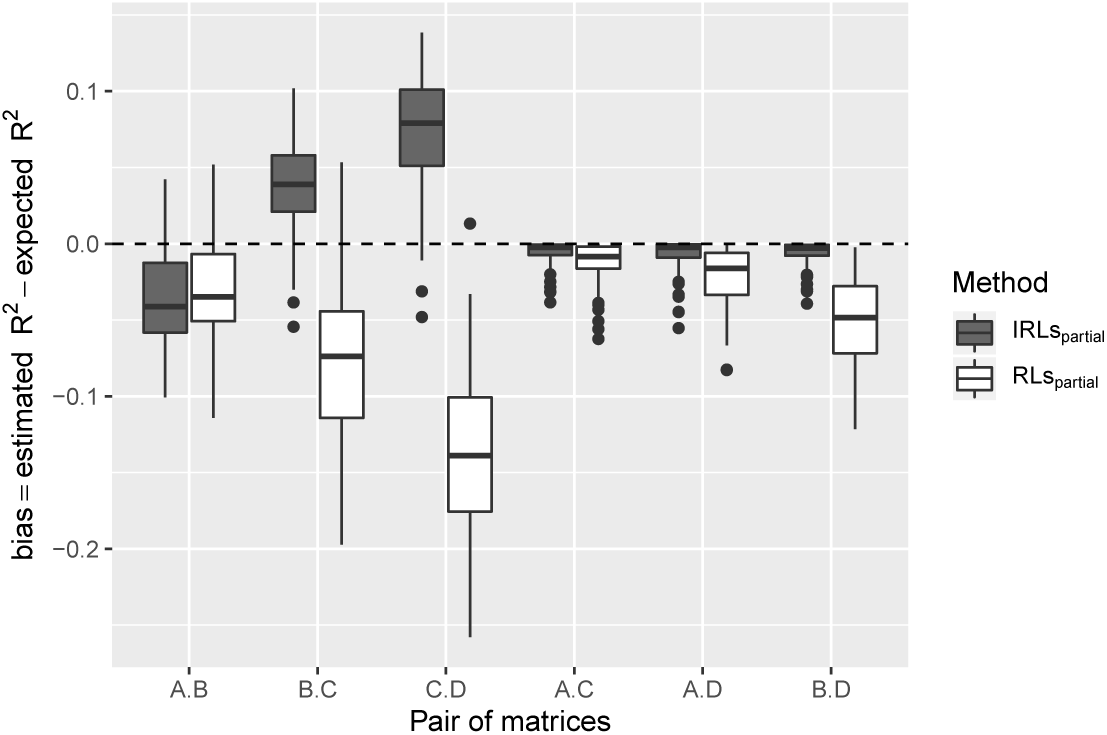
Estimation error on the partial determination coefficient. Error is defined as the absolute value of the difference between the expected and the estimated partial *R*^2^ using the corrected IRLs_*partial*_ and not corrected RLs_*partial*_ Procrustes correlation coefficient.

### 4.4 *P*_*value*_ distribution under null hyothesis

As expected *P*_*values*_ of the *CovLs* test based on the estimation of 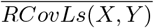 are uniformely distributed under *H*0. whatever the *p* tested (Table 2). This ensure that the probability of a *P*_*value*_ ≤ *α*-risk is equal to *α*-risk. Moreover *P*_*values*_ of the *CovLs* test are strongly linerarly correlated with those of both the other tests (*R*^2^ = 0.996 and *R*^2^ = 0.996 respectively for the correlation with `vegan::protest` and `ade4::procuste.rtest` *P*_*values*_). The slopes of the corresponding linear models are respectively 0.998 and 0.999.

**Table 2:** *P*_*values*_ of the Cramer-Von Mises test of conformity of the distribution of *P*_*values*_ correlation test to 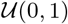 under the null hypothesis.

| p | Cramer-Von Mises p.value |  |  |
| --- | --- | --- | --- |
|  | CovLs test | protest | procuste.rtest |
| 10 | 0.203 | 0.250 | 0.194 |
| 20 | 0.682 | 0.682 | 0.687 |
| 50 | 0.560 | 0.532 | 0.551 |

### 4.5 Power of the test based on randomisation

Power of the *CovLs* test based on the estimation of 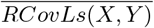 is equivalent of the power estimated for both `vegan::protest` and `ade4::procuste.rtest` tests (Table 3). As for the two other tests, power decreases when the number of variable (*p* or *q*) increases, and increase with the number of individuals and the shared variation. The advantage of the test based on the Monte-Carlo estimation of 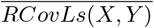 is to remove the need of running a supplementary set of permutations when IRLs is computed.

**Table 3:** Power estimation of the Procrustes tests for two low level of shared variations 5% and 10%.

| | $R^2$<br>p<br>n | 5% | | | | 10% | | | |
| --- | --- | --- | --- | --- | --- | --- | --- | --- | --- |
|  |  | 10 | 20 | 50 | 100 | 10 | 20 | 50 | 100 |
| | | $power = 1 - \beta\text{-risk}$ | | | | | | | |
| Covls.test | 10 | 0.49 | 0.45 | 0.40 | 0.45 | 0.76 | 0.68 | 0.70 | 0.68 |
|  | 15 | 0.88 | 0.80 | 0.75 | 0.75 | 0.99 | 0.98 | 0.96 | 0.95 |
|  | 20 | 0.99 | 0.96 | 0.94 | 0.93 | 1.00 | 1.00 | 1.00 | 1.00 |
|  | 25 | 1.00 | 1.00 | 0.99 | 0.98 | 1.00 | 1.00 | 1.00 | 1.00 |
| protest | 10 | 0.50 | 0.45 | 0.40 | 0.45 | 0.77 | 0.70 | 0.70 | 0.68 |
|  | 15 | 0.88 | 0.80 | 0.75 | 0.75 | 0.99 | 0.98 | 0.96 | 0.95 |
|  | 20 | 0.99 | 0.96 | 0.94 | 0.93 | 1.00 | 1.00 | 1.00 | 1.00 |
|  | 25 | 1.00 | 1.00 | 0.99 | 0.99 | 1.00 | 1.00 | 1.00 | 1.00 |
| procuste.rtest | 10 | 0.50 | 0.45 | 0.41 | 0.45 | 0.76 | 0.69 | 0.70 | 0.68 |
|  | 15 | 0.88 | 0.80 | 0.74 | 0.75 | 0.99 | 0.98 | 0.96 | 0.96 |
|  | 20 | 0.99 | 0.96 | 0.94 | 0.93 | 1.00 | 1.00 | 1.00 | 1.00 |
|  | 25 | 1.00 | 1.00 | 0.99 | 0.99 | 1.00 | 1.00 | 1.00 | 1.00 |

## 5 Discussion

Correcting the over-adjustment effect on metrics assessing the relationship between high dimension datasets has been a constant effort over the past decades. Therefore, IRLs can be considered as a continuation of the extension of the toolbox available to biologists for analyzing their omics data. The effect of the proposed correction on the classical RLs coefficient is as strong as the other ones previously proposed for other correlation coefficients measuring relationship between vector data (see Figure 3, e.g. Smilde *et al.*, 2009; SzéKely and Rizzo, 2013). When applied to univariate data, RLs is equal to the absolute value of the Pearson correlation coefficient, hence, and despite it is not the initial aim of that coefficient, IRLs can also be used to evaluate correlation between two univariate datasets. Using IRLs for such data sets is correcting for spurious correlations when the number of individual is small more efficiently than classical correction (see Figure 4, Theil, 1958).

The main advantage of IRLs over other matrix correlation coefficients is that it allows for estimating shared variation between two matrices according to the classical definition of variance partitioning used with linear models. This opens the opportunity to develop linear models to explain the variation of a high dimension dataset by a set of other high dimension data matrices.

The second advantage of IRLs is that its definition implies that the variance/co-variance matrix of a set of matrices is positive-definite. That allows for estimating partial correlation coefficients matrix by inverting the variance/co-variance matrix. The effect of the correction is less strong on such partial coefficients than on full correlation, but the partial coefficients that should theoretically be estimated to zero seem to be better identified after the correction.

## 6 Conclusion

A common approach to estimate strengh of the relationship between two variables is to estimate the part of shared variation. This single value ranging from zero to one is easy to interpret. Such value can also be computed between two sets of variable, but the estimation is more than for simple vector data subject to over estimation because the over-fitting phenomena which is amplified for high dimensional data. With IRLs and its squared value, we propose an easy to compute correlation and determination coefficient far less biased than the original Procrustean correlation coefficient. Every needed function to estimate the proposed modified version of these coefficients are included in a R package ProcMod available for download from the Comprehensive R Archive Network (CRAN).

## Acknowledgements

The authors would like to thank Dr Anthony Chariton & Dr Eric Marcon for their helpful discussions and suggestions during the writing of the manuscript.

## Appendix

### A Notations

x (vector): bold lowercase.
X (matrix): bold uppercase.
*i* = 1, …, *n*: object index.
*j* = 1, …, *p*: variable index.
*k*: iteration index.
X′: The transpose of **X**.
XY: Matrix multiplication of **X** and **Y**.
Diag(X): A column matrix composed of the diagonal elements of **X**.
X^1/2^: Matrix square root of **X**.
Trace(X): The trace of **X**.

